# CRISPRCloud2: A cloud-based platform for deconvolving CRISPR screen data

**DOI:** 10.1101/309302

**Authors:** Hyun-Hwan Jeong, Seon Young Kim, Maxime W.C. Rousseaux, Huda Y. Zoghbi, Zhandong Liu

## Abstract

The simplicity and cost-effectiveness of CRISPR technology have made high-throughput pooled screening approaches available to many. However, the large amount of sequencing data derived from these studies yields often unwieldy datasets requiring considerable bioinformatic resources to deconvolute data; a feature which is simply not accessible to many wet labs. To address these needs, we have developed a cloud-based webtool CRISPRCloud2 that provides a state-of-the-art accuracy in mapping short reads to CRISPR library, a powerful statistical test that aggregates information across multiple sgRNAs targeting the same gene, a user-friendly data visualization and query interface, as well as easy linking to other CRISPR tools and bioinformatics resources for target prioritization. CRISPRCloud2 is a one-stop shop for labs analyzing CRISPR screen data.

## Introduction

Genetic screens approaches are unbiased and rapid hypothesis generating tools allowing for the identification of novel and important biological findings. Initially, these screens were performed using chemical mutagenesis or in an arrayed manner using RNA inference (RNAi) approaches and yielded several important biological findings^1–6^. However, the advent of microarray and next-generation sequencing (NGS) technologies have rapidly moved the field towards large-scale, pooled approaches^7–11^. In earlier cases, pooled shRNA (short-hairpin RNA) libraries are barcoded and packaged into viruses, infected in a population of cells and selected for a given phenotype (e.g. growth or fluorescence). Then, hit identification was performed by hybridizing microarray^12^. In later iterations of this approach, this deconvolution step was performed using NGS^13^.

The development and optimization of CRISPR/Cas9 (Clustered Regularly Interspaced Short Palindromic Repeats and CRISPR-associated protein 9) systems have since propelled pooled screens approach to a whole new level. For instance, the robustness of hit identification has reduced the requirement for higher order redundancy in the number of targeting sgRNAs (single-guided RNAs), thus allowing for greater library diversity. Moreover, the availability of these pooled libraries on repositories such as Addgene (https://www.addgene.org/) have promoted their widespread implementation by the scientific community. Thus, the simplicity and cost-effectiveness of CRISPR technology have put pooled screens within the technical reach of most biomedical researchers^14^. While great strides that have been made on the experimental side, biologists are left with a drawback: bioinformatics complexity. To wit, the simultaneous testing of thousands of genetic perturbations demands considerable bioinformatics resources that are simply not accessible to many wet labs. Therefore, for an experimentalist to undertake such a screening project, they must first identify and collaborate with a computational biologist who is adept at analyzing such screens.

Before the CRISPR pooled screening era, several methods were proposed to ease deconvoluting data of RNAi pooled screening^15–18^, but most of those methods were not sufficient to CRISPR/Cas9 pooled screening data analysis^19^. MAGeCK (Model-based Analysis of Genome-wide CRISPR/Cas9 Knockout) was the first tool which was developed to provide CRISPR/Cas9 pooled screening data using a negative-binomial model and a modified robust ranking aggregation (RRA) algorithm^19^, and this allows deconvolution of the analysis from the beginning. HitSelect^20^, ScreenBEAM (Screening Bayesian Evaluation and Analysis Method)^21^, BAGEL (Bayesian Analysis of Gene Essentiality)^22^, sgRSEA (single-guide RNA Set Enrichment Analysis)^23^, PBNPA (Permutation based non-parametric analysis of CRISPR/Cas9 screen data)^24^, and MAGeCK-VISPR^25^ proposed to provide a more accurate deconvolution of the data with different statistical models. However, those analysis tools tend to be script-based because they were developed for bioinformaticians or scientists who are very computationally savvy. Of the newly-developed tools, the most user-friendly are CRISPRAnalyzeR^26^, CRISPRcloud^27^, and PinAPL-Py^28^, as they have web-based interfaces and represents as a first-step toward enabling scientists who are actually generating the CRISPR7/Cas9 screen data to analyze these large dataset. However, each of these tools have various of rate-limiting steps such as requiring intricate tuning of parameters for trimming and mapping data, long transfer times and file copying errors in the transfer of a large amount of sequence data over the internet, lack of fast and powerful statistical tools etc. Overall, the promise of an online tool for researchers is still unfulfilled.

In light of those challenges, we have developed CRISPRCloud2 (CC2, http://crispr.nrihub.org). CC2 is a one-stop shop for researchers with zero programming background to pre-process, perform quality checks, apply statistical analyses, query, and visualize their data (**Table 1**).

**Table 1.** Feature table of CRISPRCloud2 as compared with existing methods. HitSelect, ScreamBEAM, PBNPA, and sgRSEA are not listed in this table because they are script-based and do not provide the non-statistical features listed below.

| Features | CRISPRCloud2 | caRpools<sup>29</sup> | CRISPRAnalyzeR<sup>26</sup> | MAGeCK-VISPR<sup>25</sup> | PinAPL-Py<sup>28</sup> |
| --- | --- | --- | --- | --- | --- |
| Web interface | ✓ |  | ✓ |  | ✓ |
| Cloud service | ✓ |  |  |  |  |
| Parameter free trimming | ✓ |  |  | ✓ |  |
| Installation free | ✓ |  | ✓ |  | ✓ |
| Web-based FASTQ file processing | ✓ |  | ✓ |  | ✓ |
| Automatic reverse complement FASTQ file detection | ✓ |  |  | ✓ |  |
| Gene-level statistics | ✓ | ✓ | ✓ | ✓ | ✓ |
| sgRNA-level statistics | ✓ | ✓ | ✓ | ✓ | ✓ |
| Multiple group studies | ✓ | ✓ | ✓ | ✓ | ✓ |
| Quality control | ✓ | ✓ | ✓ | ✓ | ✓ |
| sgRNA mapping algorithm | ✓ | ✓ | ✓ | ✓ | ✓ |

## Results

### Architecture of CRISPRCloud2

Building web-based analysis platform for big data analysis is a challenging task. The first challenge lies in that large files has to be transferred over the Internet. Raw FASTQ files from CRISPR/Cas9 pooled screening can be substantial (about 1~10 GB per a sample) and sending those large files through the Internet is not practical. Indeed, other web-based platforms which require uploading FASTQ files are having an issue with the large size datasets^30^. Furthermore, and perhaps more importantly, uploading raw data has data-privacy implications, which is becoming a major concern recently^31^.

To overcome these challenges, we developed CC2 using the Amazon Web Service (AWS) environment (**Fig. 1**). It is compatible with most modern web browsers. The fast client-side gRNA mapping program implemented can reduce input files of several gigabytes into a single megabyte-size file. By transferring the much smaller count file through the Internet, CC2 decreases the transfer time and prevents the sharing of raw input files. Our adaptive mapping algorithm implemented using Angular (https://angular.io/) and TypeScript (https://www.typescriptlang.org) provides an open-source front-end web application platform.

**Figure 1.**
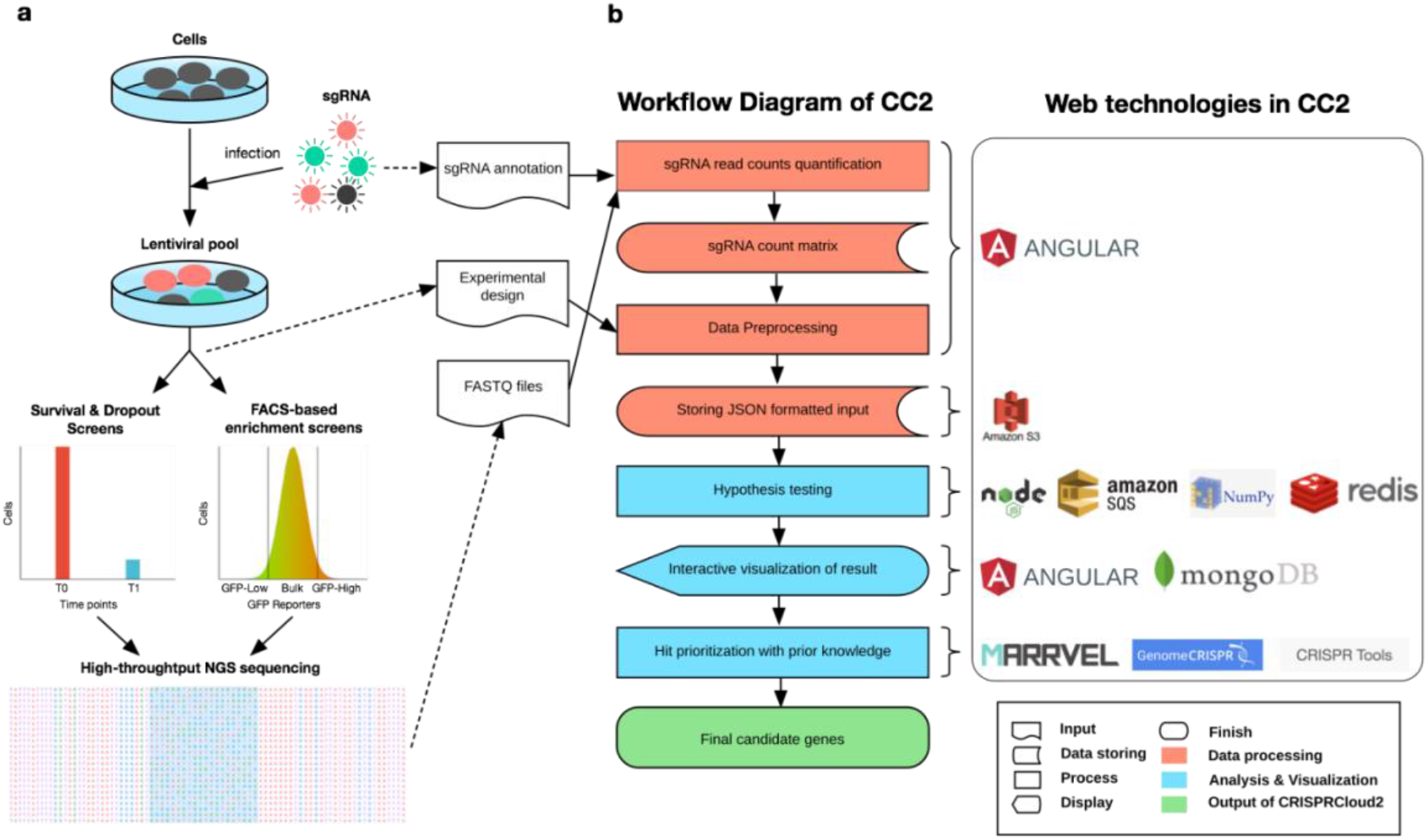
The illustration of CRISPR/Cas9 pooled screening experiment, workflow diagram, and configuration for CRISPRCloud2. (a) The schematic illustration of a CRISPR/Cas9 pooled screening experiment. Each dotted arrow line links between an experimental step and an input of CRISPRCloud2. (b) The workflow of CRISPRCloud2. This entire workflow runs on Amazon web services (AWS), and various web-based technologies were used to build this web-based cloud computing platform.

Another challenge is the huge demand on computing power. Platforms built with a centralized server solution will have load-balancing problems when many users are submitting their requests simultaneously, resulting a much longer user waiting time and even system-wide failure. To address this challenge, CC2 provides a decentralized and a cloud-computing based scalable service through the combination of AWS infrastructure (**Fig. 1**) including Amazon Elastic Compute Cloud (EC2) (https://aws.amazon.com/ec2/), Amazon Simple Storage Service (S3) (https://aws.amazon.com/s3/) and Amazon Simple Queue Service (SQS) (https://aws.amazon.com/sqs/).

### The adaptive hash-mapping algorithm provides a fast and accurate alignment

To map CRISPR/Cas9 screen data to a reference library accurately, we introduced an adaptive hash-mapping algorithm that is both fast and extremely accurate. We tested the algorithm on five published datasets (**Supplementary Table 1**), and our results demonstrate better mappability than MAGeCK and PinAPL-Py (**Fig. 2**), at comparable speeds. Our scalable cloud-based architecture, coupled with the binary presentation-based algorithm, processes millions of reads in a matter of minutes. CC2 is a few seconds slower than MAGeCK which is a stand-alone application, but it is more accurate than other methods. CC2 is currently the only CRISPR/Cas9 online screen analysis tool with parameter-free guide RNA (gRNA) level quantification.

**Figure 2.**
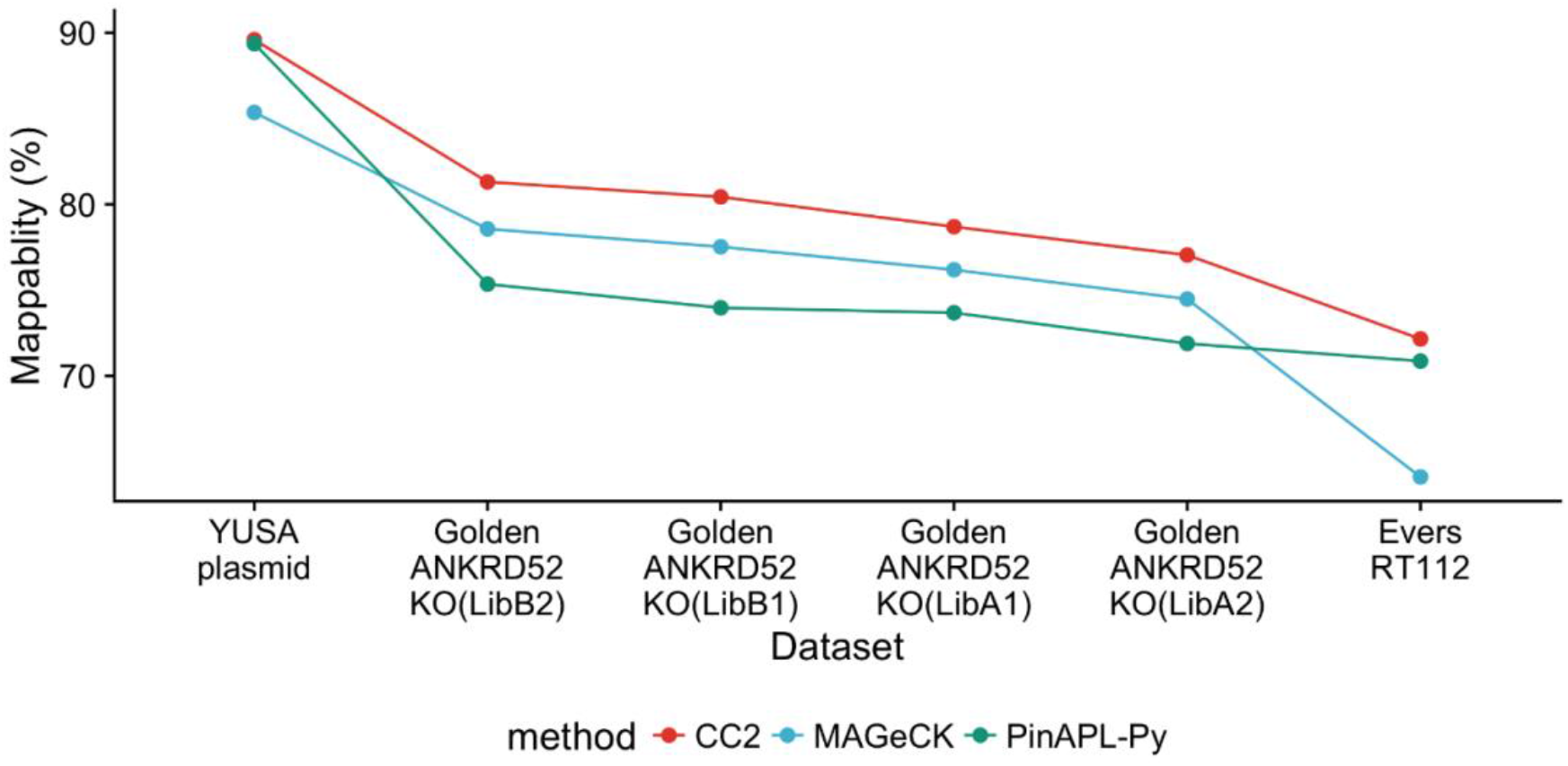
CRISPRCloud2 outperforms MAGeCK and PinAPL-Py in the percentage of mapped reads in six benchmark datasets.

To understand the performance differences, we studied the reads that are mapped by CC2 but not MAGeCK or PinAPL-Py (**Fig. 3**). We observed that only 64% of the reads were mapped by MAGeCK compared to CC2 in Evers RT112 dataset (**Fig. 2**). This is primarily due to the fact that MAGeCK estimates the trimming window using the first N reads from the input (N is 100,000 by default). There is no guarantee that these windows are optimal for the rest of the input files (**Fig. 3b**). PinAPL-Py uses cutadapt^32^ for the read trimming and bowtie2^33^ for the mapping using the local alignment mode. We observed that the local alignment of bowtie2 failed due to the incorrect trimming from cutadapt (**Fig. 3b**).

**Figure 3.**
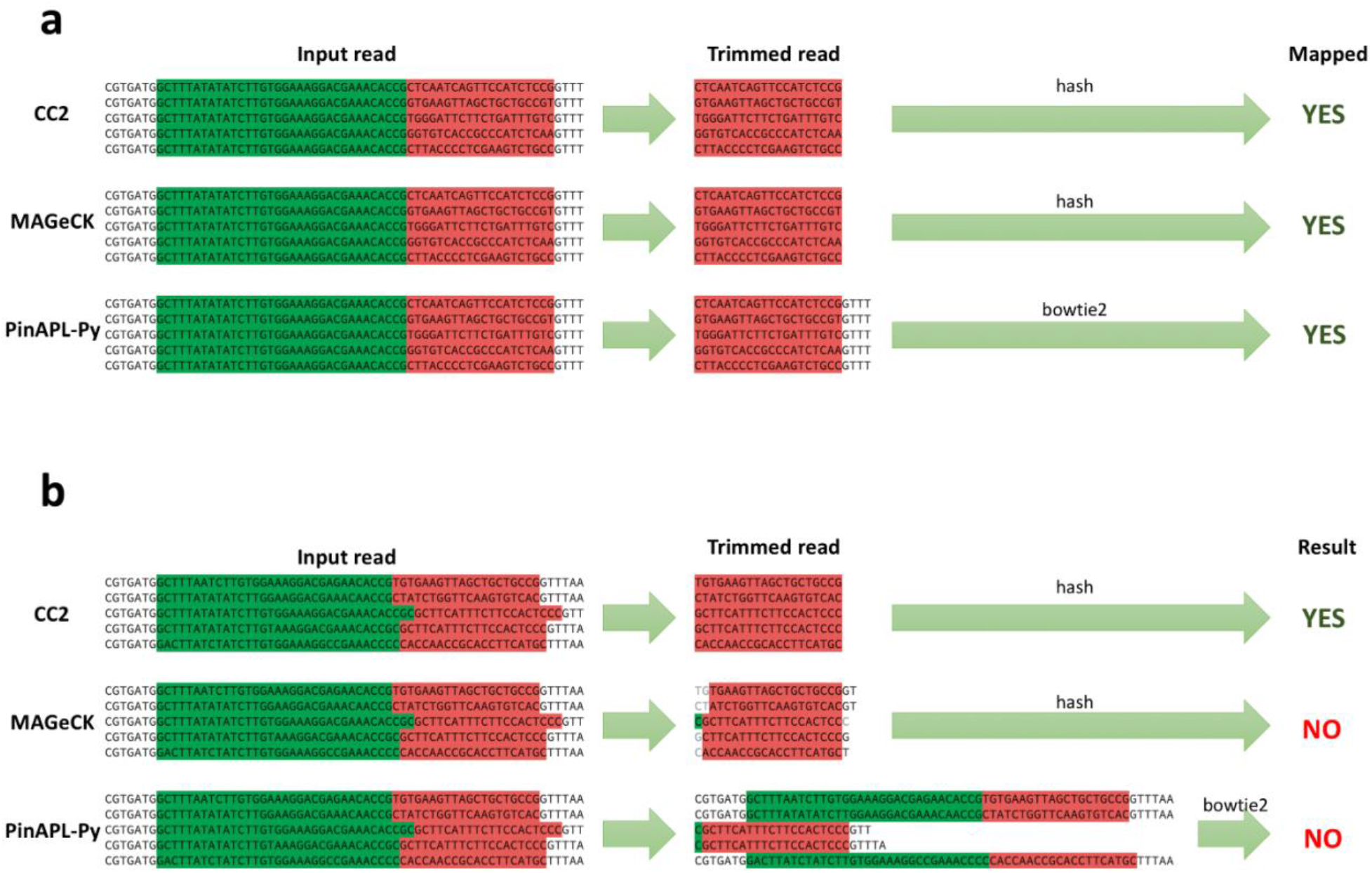
Examples of mapping results of CC2, MAGeCK, and PinAPL-Py. Examples are taken from Evers et al.’s CRISPR RT112 cell line screening data. (a) All three methods successfully detect sgRNA sequences. (b) Only CC2 successfully identifies sgRNA sequences while MAGeCK and PinAPL-Py failed to identify the correct sgRNA sequences from the reference library.

### CRISPRCloud2 offers robust target identification

Identifying candidates by statistical hypothesis-testing is the second key component in any screen analysis. In CC2, we have adopted a beta-binomial model^34^ with a modified Student’s t-test to measure differences in single-guide RNA (sgRNA) levels, followed by Fisher’s combined probability test to estimate the gene level significance. In this regard, CC2 is five hundred times faster than CRISPRcloud for a genome-wide CRISPR/Cas9 screening dataset (**Supplementary Table 2**). To evaluate the statistical power of CC2, we compared it with six state-of-the-art methods on three benchmark datasets evaluating gene essentiality^35^ using CRISPRn (CRISPR nuclease gene knockout via Cas9and CRISPRi (a CRISPR/Cas9 system with an inactive Cas9 fused to the transcriptional repressor KRAB which results in gene repression) technologies (**Table 2**). These benchmark datasets were constructed based on 46 genes that are essential for cell survival and 47 genes that are non-essential. As shown in **Fig. 4** and Supplementary **Fig. 1** and 2, CC2 outperforms all other methods at every FDR cut-off level. All methods demonstrated a small type-I error due to the strong lethality phenotype of the CRISPR assay, but CC2 demonstrated a significantly lower type-II error than all the other methods (**Supplementary Fig. 2**). Furthermore, we also find that other methods are not able to detect some of the essential genes (i.e. COPS8, RPL5, and RPL27, except COPS8 and RPL5 in CRISPRi-RT112 screening) even when more than half of sgRNAs for these genes showed differential negative abundance between the two-time points (**Fig. 5 and Supplementary Figs. 3 and 4**). Across datasets from two cell lines and two CRISPR/CRISPRi libraries, with false discovery rates (FDR) ranging from 10% to 0.01%, CC2 had a much larger F1-score and recall. CC2 is thus both accurate and sensitive.

**Figure 4.**
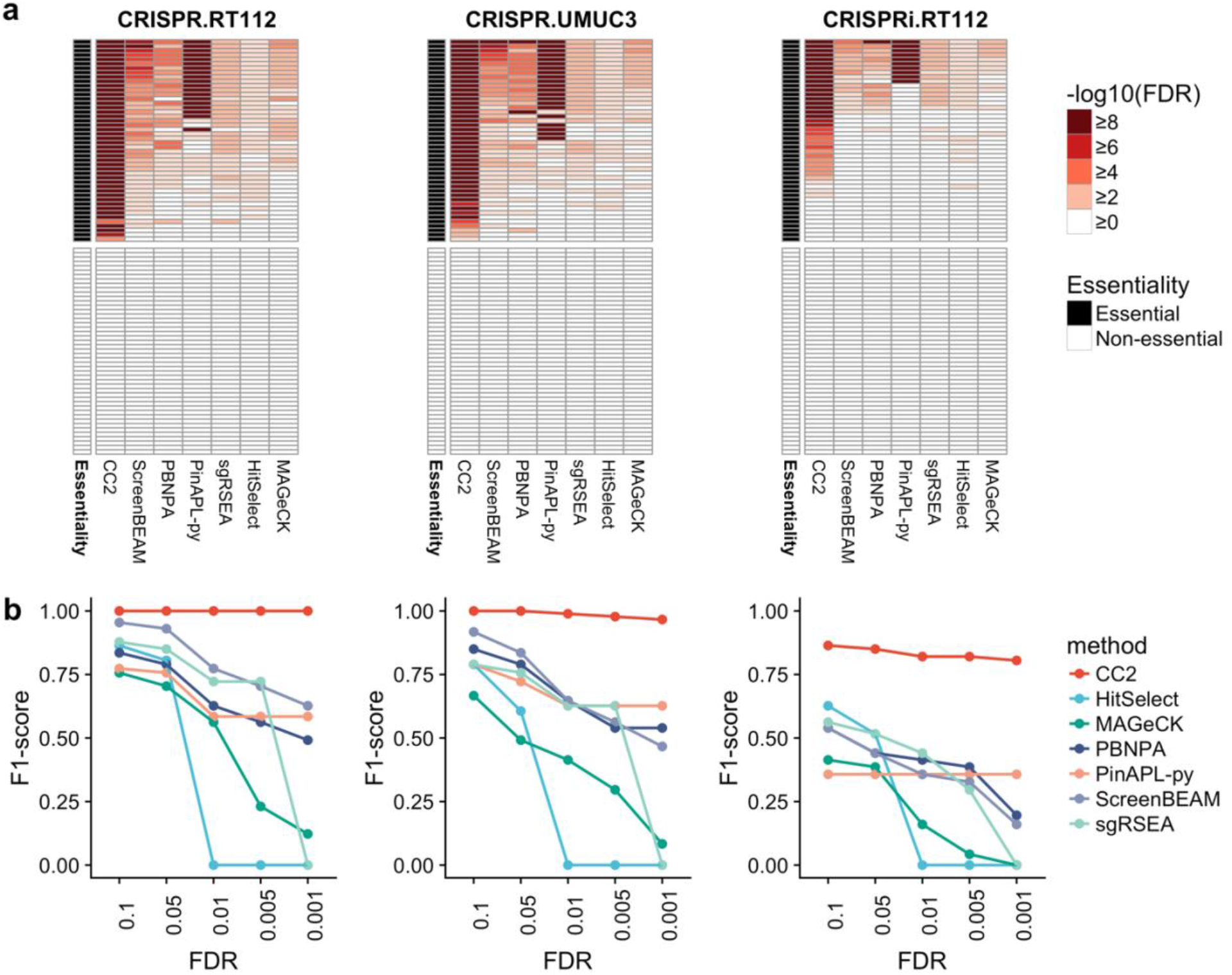
CC2 offers robust target identification with high precision and recall. (a) Heatmaps illustrate FDRs of gene statistics from each of six leading high-complexity pooled screen analysis tools. The color of the cells indicates gene essentiality (black color for essential genes, white color for nonessential genes). (b) F1-score measurements at different FDR cut-offs across all methods. At various commonly used FDR cut-off, CC2 was able to identify most of the essential genes with high precision and recall rate.

**Figure 5.**
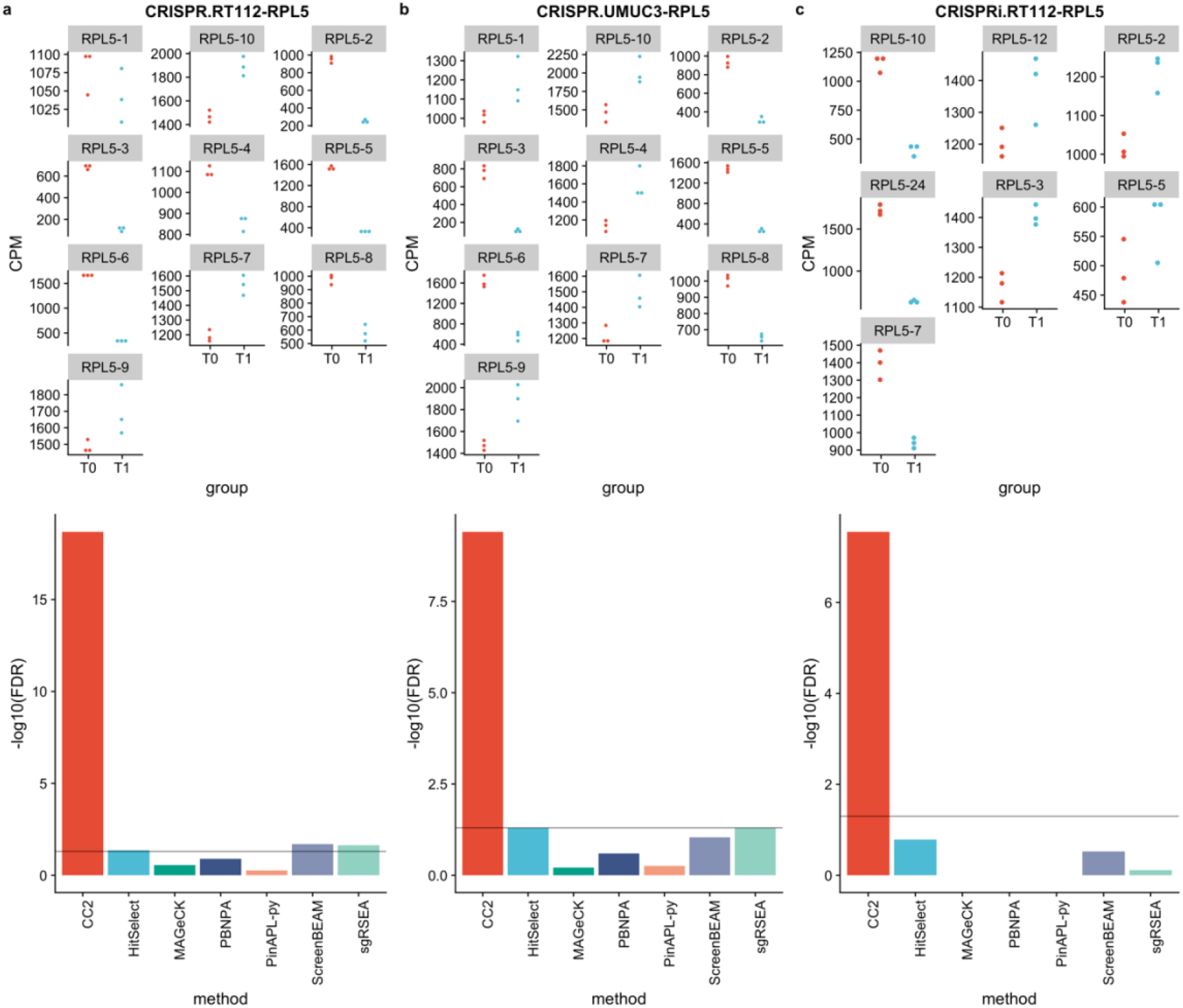
CRISPRCloud2 detects essential genes missed by other leading methods: the case of RPL5. sgRNA quantification for RPL5 in cell line RT112 (a), UMUC3 (b) using CRISPR and RT112(c) using the CRISPRi library. The FDR value for RPL5 in each screen is plotted across all the methods. A horizontal line at FDR=0.01 is used as a threshold for statistical cutoff. CC2 outperforms all other methods of identifying RPL5 as an essential gene across all benchmark datasets.

**Table 2.**
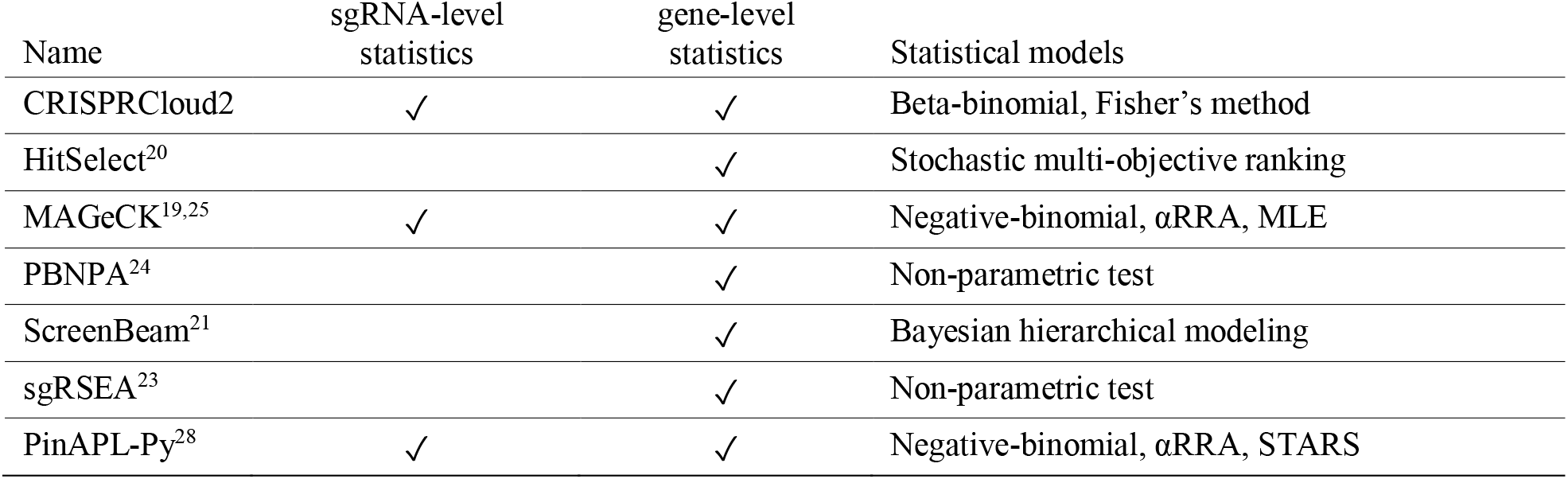
Statistical models used by CRISPRCloud2 and existing methods. All of the methods were used in the target identification benchmarking.

## Discussion

The advent of CRISPR-Cas9 systems as powerful and refined genome manipulation techniques has heralded a new era of large-scale screening approaches. Indeed, over the past three years, there has been exponential growth in the number of pooled genetic screens. The number of datasets for CRISPR/Cas9 screens in Gene Expression Omnibus have more than tripled each year (39 datasets in 2015, 121 datasets in 2016, and 408 datasets in 2017). Much of this has been due to the widespread availability of large-scale genome-wide perturbation libraries via the non-profit repository Addgene (https://www.addgene.org/) and resource sharing between labs. Cheaper and faster sequencing options have also improved the accessibility of this type of approach to virtually any lab. However, while the major limitations of these first two aspects have been lifted, the computational toll that data deconvolution takes has rendered these approaches daunting to many. In this study, we lifted this last barrier by providing a framework – CRISPRCloud2 – that is straightforward, multifunctional and does not require heavy computation. This pipeline is a one-stop-shop for individuals performing CRISPR-based screens and outperforms, to the best of our knowledge, all other current CRISPR-screen deconvolution programs.

Because we intend CC2 to be accessible to researchers with no background in programming, we designed the interface to guide the user through each step of the analysis, from uploading raw data to selecting target hits based on summary statistics. We have also developed a suite of cloud-based solutions for data query and visualization (**Supplementary Methods**). Moreover, we complement the site with a video tutorial as well as a step-by-step guide to running one’s data through the pipeline. Importantly, results obtained from CC2 (i.e. “hits” identified in the screen) can then be immediately linked to other CRISPR/Cas9 screen analysis tools such as GenomeCRISPR^36^, CRISPRTools^37^, as well as MARRVEL^38^. The latter approach allows the user to query the identified hits for their human disease relevance and functional annotation in model organisms, thus promoting functional prioritization. With the increasing applications of CRISPR/Cas9 systems proliferating, particularly in regard to multiplexing/pooled approaches, the need for accessible bioinformatics tools grows more pressing. With CC2, the research community has a robust and user-friendly online tool to mine this data with ease.

## Methods

### Statistical hypothesis testing using beta-binomial distribution for sgRNA-level differential analysis

We adopted a beta-binomial model proposed for Serial Analysis of Gene Expression (SAGE) by Baggerly *et al*.^34^. Specifically, let *p_i_* be the true proportion of an sgRNA in sample *i*. We assume the value of *p_i_* can vary from sample to sample and follows a beta distribution, *p_i_*~*Beta*(*α,β*). Let *X_i_* denote the number of read counts for a sgRNA in the *i^th^* sample. We assume *X_i_* follows a binomial distribution, *X_i_*|*p_i_*~*Binomial*(*n_i_, p_i_*), where *n_i_* is the total number of mapped reads in sample *i*. To combine the estimated 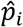 across multiple samples of the same treatment group, a linear model is proposed *p^A^* = Σ*w_i_p_i_*, where *i* is the index for samples and *w* is the weight vector for samples in group A. Baggerly *et al*. proved that as long as *w^T^* 1 = 1, the expectation of *E*(*p^A^*) is unbiased. The value of *w* is estimated through gradient descent methods by minimizing the variance on *p^A^*. Baggerly *et al*. showed that 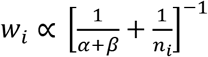.

CRISPRCloud2 performs the sgRNA-level differential analysis between two groups using a Student *t*-test like statistic proposed by Baggerly *et al*.:

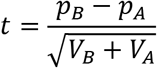

where *p_A_* and *p_B_* are the proportion of sgRNA, and *V_A_* and V_B_ are the group variance of sgRNA, for groups A and B, respectively. Test statistic *t* represents the strength of the difference of sgRNA abundance between groups *A* and *B*. In other words, a large positive *t*-value indicates that the quantity of sgRNA in group *B* is more than in group *A*, and a large negative *t*-value indicates that the quantity of sgRNA in group *B* is less than in group *A*.

The variance is estimated by

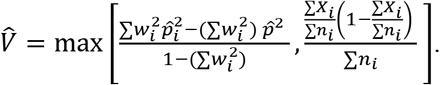

To measure the statistical significance of the difference, we approximate the *p*-value of a given *t* in a Student’s *t*-distribution with a degree of freedom (*df*) defined by

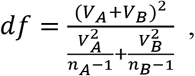

where *n_A_* and *n_B_* are the numbers of replicates in groups *A* and *B*.

A sgRNA log_2_ fold-change in abundance between A and B (log_2_ *FC*) is estimated by

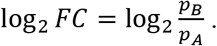

#### sgRNA p-value aggregation for gene-level statistics

Because multiple significant sgRNAs targeting the same gene hold greater biological significance than a single significant sgRNA, we must aggregate *p*-values to increase confidence in target identification. To do so, we combine *p*-values of sgRNAs for a target gene using Fisher’s method^39^ to assess overall differences at the gene level. The combined chi-square statistical test is used:

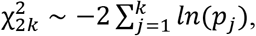

where *k* is the number of sgRNAs targeting a gene in the screen and *p_j_* is the *p*-value of *j*-th sgRNA for the gene. *χ*^2^ follows a chi-squared distribution with 2*k* degrees of freedom. To correct for multiple hypothesis testing, we adopted the Benjamini-Hochberg procedure^40^ to estimate the False Discovery Rate (FDR).

### Gene-level statistics benchmarking on existing methods

We used three different CRISPR/CRISPRi pooled screen datasets from Evers *et al*. ^11^(RT112 and UMUC3 cell line screens with CRISPR; RT112 cell line screen with CRISPRi), which provide ground-truth labels of essentiality for each gene. With those screening datasets and labels, we benchmarked the accuracy of essential gene detection by CC2 with six other published methods (**Table 2**). We computed the False Discovery Rate (FDR) for each gene from each method in the benchmark and set five different levels of FDR cut-off (0.1, 0.05, 0.01, 0.005, 0.001) for essential gene classification. For example, if we set FDR cut-off to 0.1, then a gene is predicted to be essential in the cell line if the FDR of the gene falls below the cut-off value. We calculated recall (a recall value close to 1 indicates a prediction with a low false negative rate), precision (a precision value close to 1 indicates a prediction with a low false positive rate), and F-measure (the harmonic mean of precision and recall) of all the methods at each FDR level. All of the data and scripts for the benchmarking are available at https://github.com/hyunhwaj/CC2-bench. Parameters used in these experiments are described below.

#### CRISPRCloud2

Benchmarking of CRISPRCloud2 was performed without parameter tuning since CC2 is parameter-free. FDR values for negative changes between two different time points (T0 and T1) from CC2 statistical analysis were used for benchmarking.

#### MAGeCK

MAGeCK version 0.5.6 was used for benchmarking. We ran MAGeCK with the ‘mageck test’ command with the following parameters: --norm-method and --adjust-method. We performed 100 permutations for the modified robust ranking aggregation (RRA) algorithm to estimate the gene-level statistics on the benchmark datasets.

#### ScreenBEAM

ScreenBEAM R package (version 1.0.0, https://github.com/jyyu/ScreenBEAM) was used for benchmarking ‘data.type’ parameter was set as ‘NGS,’ and ‘do.normalization’ was set as TRUE, ‘nitt’ and ‘burnin’ parameters for Bayesian computing were set at 15000 and 5000. ScreenBEAM does not provide the one-sided *p*-value for negative selection, so for the FDR comparison with other methods, we changed the FDR of a gene to 1 if the *β* of the gene is greater than 0.

#### sgRSEA

sgRSEA R package (version 0.1, https://cran.r-project.org/web/packages/sgRSEA/) was used for benchmarking. we set the multiplier at 30.

#### PBNPA

PBNPA R PACKAGE (version 0.0.2, https://cran.r-project.org/web/packages/PBNPA/) was used for benchmarking. We set the sim.no parameter at 10.

#### HitSelect

We ran HitSelect MATLAB package (https://github.com/diazlab/HiTSelect). Normalization by Sequencing depth option was selected for benchmarking.

#### PinAPL-py

We used the PinAPL-py website (http://pinapl-py.ucsd.edu) to perform the benchmarking. For the sgRNA read counting, we used ‘GGCTTTATATATCTTGTGGAAAGGACGAAACACCG, GCTTTATATATCTTGTGGAAAGGACGAAACACCG,’ and ‘CTTTATATATCTTGTGGAAAGGACGAAACACCG,’ were used for ‘seq_5_end’ parameters of ‘CRISPR-RT112’, ‘CRISPR-UMUC3’, and ‘CRISPRi-RT112’ datasets. We used CPM normalization and set the GeneMetric parameter as ‘aRRA’ to perform a modified robust ranking aggregation (RRA). We used the combined FDR values for each gene in the benchmarking.

### Algorithm for quantifying sgRNA abundance

#### Previous methods and limitations of CRISPRCloud1

Recently published tools for CRISPR pooled screen analysis, including CRISPRcloud (CC1)^27^, MAGeCK^25^, CarRpools^29^, CRISPRAnalyzeR^26^, and PinAPL-Py^28^, provide different methods for estimating the abundance of sgRNAs in each sample from pooled libraries. In most cases, input data consist of raw FASTQ-format sequencing result files. CC1 is the first tool to offer an online user-defined, light-weight quantification method which proceeds on the user-client side. In contrast, CRISPRAnalyzeR and PinAPL-Py run their quantification methods on the server-side. As a result, CC1 minimizes information passed through the Internet by transferring only the processed count matrix to the cloud storage. However, the quantification algorithm of CC1 is limited in that it could have a reduced sgRNA coverage if the location patterns of sgRNAs in raw-sequencing files is ‘staggered,’ as pointed out by Spahn *et al*.^28^. This problem happens because CC1 extracts sgRNA sequence for each read at a fixed location^28^. Another limitation of CC1 is the fact that the user must decide where the extraction site is. Nevertheless, CC1 does not require tuning and is thus arguably more user-friendly than other tools. For instance, in CRISPRAnlyzer and PinAPL-Py, users are required to set many tuning parameters for sgRNA quantification, such as adapter sequence, sgRNA sequence length, and whether sgRNA sequence reads are reverse-complement. Improperly setting these mandatory parameters can hinder non-bioinformaticians from using these tools.

In CC2, we performed extensive software engineering to address these issues. As a result, users no longer need to perform complicated parameter tuning for the sgRNA abundance quantification; one must simply provide the input files to CC2.

#### The binary representation of sgRNA sequence lowers the cost of computation

We used a binary representation for sgRNA sequence. This approach is memory-efficient and improves the user experience at the client-side^41^. It only needs *max*(*2K, M*) bits to store an sgRNA-sequence, where *M* is the length of the sequence and *K* is the bit size to store a primitive integer in the machine (usually 64 bits) because we only need two bits to save a nucleotide (i.e., ‘A’ is ‘00’, ‘C’ is ‘01’, ‘G’ is ‘11’, and ‘T’ is ‘10’). The memory size is about half of that required for storing a character string of the sequence, i.e., 160 bits are needed to store a 20nt sgRNA sequence. Another benefit of binary representation is that it lowers the time complexity for the shift operator when comparing all k-mers of an sgRNA read using a sliding window. This is an essential function for the quantification algorithm in CC2. Compared to the string shift operator functions, such as string copy, substring extraction, and concatenation, the binary representation produces dramatically shorter running times.

#### Sliding window-based algorithm gives a high-resolution quantification with comparable running time

With a binary representation, we run the quantification algorithm as follows: First, we build a hash table for the reference library, with each key of the library in the hash table converted to the binary representation. Second, for each read, we scan the sequence of the read from 5′ to 3′ with the sliding window. In the *i*-th iteration, the sliding window contains a substring of the read sequence from *i* to *i* + *k* – 1, where *k* is the length of the sgRNAs. The substring is also converted to a binary sequence, and the hash table is quickly checked to see if the sequence in the sliding window exists in the reference library. If the sequence is found in the hash table, then the count of the sequence is increased by one, and then the algorithm proceeds to the next read. Otherwise, it moves to *i* + 1-th iteration and the bit-shift method will be applied to take the next sliding window.

For the case of a reverse complement sequenced sample, the entire procedure is repeated on the reverse complement reference sgRNA library and scanning the read from 3′ to 5′. After both assays are performed (5′ to 3′ and 3′ to 5′ with the reverse complement reference sequences), mapping results between both sequences are compared. The one with a larger count corresponds to the correct sequence mapping. We compared the mappability and running time of CC2 to those of MAGeCK^19^ and PinAPL-Py^28^ across multiple datasets from previous studies (**Figure 2**). To perform a judicious comparison, we re-implemented every algorithm in javascript, except the cutadapt adaptive trimming of PinAPL-Py. While not the fastest of the test cohort, the processing speed of CC2 is comparable to that of MAGeCK.

## Acknowledgements

This work has been supported by National Institute of General Medical Sciences R01-GM120033, National Science Foundation - Division of Mathematical Sciences DMS-1263932, Cancer Prevention Research Institute of Texas RP170387, Houston Endowment (Z.L.), Huffington Foundation, Howard Hughes Medical Institute (H.Y.Z.), and the Parkinson’s Foundation Stanley Fahn Junior Faculty Award PF-JFA-1762 (M.W.C.R.).

## Author contributions

H-H.J., M.W.C.R., and Z.L. designed the study. H-H.J. and S.Y.K. implemented the CRISPRCloud2 software. H-H.J. and Z.L. performed analysis. H.Y.Z. and Z.L. supervised the project. H-H.J., M.W.C.R., and Z.L. wrote the manuscript with input from all the authors.

